# Active and repressed chromatin domains exhibit distinct nucleosome segregation during DNA replication

**DOI:** 10.1101/418707

**Authors:** Thelma M. Escobar, Ozgur Oksuz, Nicolas Descostes, Roberto Bonasio, Danny Reinberg

## Abstract

Chromatin domains and their associated structures must be faithfully inherited through cellular division to maintain cellular identity. Yet, accessing the localized strategies preserving chromatin domain inheritance, specifically the transfer of parental, pre-existing nucleosomes with their associated post-translational modifications (PTMs) during DNA replication is challenging in living cells. We devised an inducible, proximity-dependent labeling system to irreversibly mark replication-dependent H3.1 and H3.2 histone-containing nucleosomes at single desired loci in mouse embryonic stem cells such that their fate after DNA replication could be followed. Strikingly, repressed chromatin domains are preserved through the local re-deposition of parental nucleosomes. In contrast, nucleosomes decorating active chromatin domains do not exhibit such preservation. Notably, altering cell fate leads to an adjustment in the positional inheritance of parental nucleosomes that reflects the corresponding changes in chromatin structure. These findings point to important mechanisms that contribute to parental nucleosome segregation to preserve cellular identity.

## Introduction

Genome function and cellular identity are maintained through the structural organization of chromatin domains that either facilitate or impede transcription. The presence of specific histone post-translational modifications (PTMs) within these chromatin domains not only correlate with a given transcription status, but also facilitate the formation of repressive chromatin structures that impact gene expression (Reinberg and Vales, 2018). In order to maintain the integrity of these gene expression programs through cell division, the molecular features defining these chromatin structures must be transmitted, rather than established anew. Important determinants to establishing different types of chromatin states are the histone PTMs decorating primarily the tails of histone H3 and H4 (Stillman, 2018). Thus, in addition to DNA replication, chromosome duplication should entail the accurate reassembly/segregation of parental nucleosomes onto the appropriate locale of each daughter DNA molecule. This latter process involves a tightly coupled deposition of histones to the replication machinery behind the replication fork (Cusick et al., 1981; Herman et al., 1981). The founding studies on the structure of replicated chromatin establishes that parental histones are deposited onto newly synthesized DNA relatively quickly and that both replicated DNA strands capture equal amounts of parental histones (Annunziato, 2013; Petryk et al., 2018; Yu et al., 2018). It is now accepted that parental canonical histones, *i.e.* replication-dependent histones, rapidly re-assemble behind the replication fork starting with the H3-H4 tetrameric core followed by H2A-H2B dimer deposition (Campos and Reinberg, 2009; Xu et al., 2010; Alabert and Groth, 2012; Campos et al., 2014). Moreover, clear evidence that specific histone PTMs have the potential to be transmitted across mitosis (epigenetically inherited) was derived from studies of two distinct methylated states of histone H3 within the (H3-H4)_2_ core: tri-methylated H3K9 (H3K9me3) and di- or tri-methylated H3K27 (H3K27me2/me3) (Grewal and Moazed, 2003; Margueron et al., 2009; Margueron and Reinberg, 2011) and a model to account for this inheritance was recently put forward (Reinberg and Vales, 2018). Notably, both of these histone modifications are associated with repressed chromatin domains.

Nucleosomes vary in the composition of their histone components. While only one histone H4 isoform has been identified, there are several histone H3 variants including H3.1 and H3.2 that differ at one amino acid and are considered the canonical replication-coupled versions (Tagami et al., 2004). The observation that parental H3.1- and H3.2-containing nucleosomes are re-deposited as intact (H3.1-H4)_2_ or (H3.2-H4)_2_ tetramers upon DNA replication (Xu et al., 2010), supports a model for the local inheritance of histone PTMs. However, direct testing for the local re-deposition of parental canonical tetramers at a particular locus has not been achieved. In particular, the *in vivo* re-deposition of parental histones within the general vicinity of their original genomic position has not yet been examined through direct methods, but instead through proteomics, marking of newly replicated DNA and ChIP-sequencing techniques (Zee et al., 2012; Alabert et al., 2014; Clement et al., 2018; Reveron-Gomez et al., 2018). Although, the combination of these approaches provides insights into the “bulk” re-deposition of parental nucleosomes, these studies cannot determine the fidelity of such re-deposition at a given chromatin domain, key to tackling the mechanisms of epigenetic inheritance.

In the case of embryonic stem cells (ESCs), their pluripotent state depends on both the active transcription of genes encoding the ‘pluripotent factors’, *i.e. Pou5f1*, *Nanog* and *Sox2* (among others), and the repression of genes encoding lineage-specifying developmental regulators (Boyer et al., 2006; Ng and Surani, 2011; Dowen et al., 2014). The Polycomb group (PcG) of proteins maintain the status of such gene repression in part through: 1) the catalysis of H3K27me2/me3 by PRC2 and 2) H3K27me2/me3 providing a platform for the recruitment of PRC1 and chromatin compaction (Boyer et al., 2006; Margueron and Reinberg, 2011; Pengelly et al., 2013; Simon and Kingston, 2013), although alternative pathways have been proposed (Cooper et al., 2016; Almeida et al., 2017). Indeed, perturbations of PcG proteins can aberrantly affect differentiation and foster a loss in cellular identity (Boyer et al., 2006; Margueron and Reinberg, 2011). Thus, the integrity of chromatin domains must be conveyed in a precise spatial and temporal manner to ensure proper development and to establish the maintenance of cellular identity.

To gain insights into how parental nucleosome re-deposition might contribute to epigenetic inheritance, we devised a method that permanently and specifically labels H3.1- and H3.2-containing nucleosomes at desired actively transcribed or repressed genes in ESCs. This system allows us to follow the fate of the “marked” parental nucleosome upon DNA replication when ESCs are in a pluripotent state or enter into a specific differentiation program. The findings reveal that nucleosomes from repressed (but not active) chromatin domains are re-deposited within the same chromatin domain after DNA replication, whereas parental nucleosomes from actively transcribed genes are dispersed. Moreover, by inducing an altered cell fate from ESCs, the local re-deposition of parental nucleosomes during DNA replication at defined loci now reflects their conversion from a repressed to an active chromatin domain. These findings point to the inheritance of repressed versus active chromatin domains.

## Results

### Marking nucleosomes at a specific locus to follow their segregation upon DNA replication

To investigate the segregation of parental nucleosomes, we developed a bio-orthogonal system to irreversibly mark H3.1 and H3.2 *in vivo* at candidate loci to follow their re-deposition at the single nucleosome level across cellular division in mouse ESCs (Figure 1A). First, we introduced a Biotin Acceptor Peptide (BAP) motif sequence into the N-terminus of the endogenous H3.1 and H3.2 loci to biotinylate H3 chromatin using the *Escherichia coli* Biotin Ligase (BirA) (Kulyyassov et al., 2011; Shoaib et al., 2013). To recruit BirA to biotinylate nucleosomes at specific loci on chromatin, BirA was fused to the catalytically inactive dCas9. To ensure the exclusive labeling of parental histones, we restricted biotinylation to a tight temporal window by expressing dCas9-BirA from an integrated cassette under the control of a doxycycline-inducible promoter and incorporating an FKBP degradation domain (DD) (Banaszynski et al., 2006). With this system in hand, the expression of chosen gRNAs allowed us to control spatial (gene) specificity of biotinylation (Chen et al., 2013), resulting in the desired, precise deposition of the biotin tag, exclusively at the chosen locus of interest (Figure 1A). We then created clonal dCas9-DD-BirA-expressing ESCs containing Flag-BAP knock-ins to the N-terminus of 13 endogenous copies of replication-dependent H3.1 and H3.2 in the *Hist1h3* cluster (Figures 1A and S1A-B). Chromatin immunoprecipitation (ChIP) followed by western blots of Flag-BAP-H3 showed that the tagged-histones were incorporated into both active and repressive chromatin as defined by the presence of H3K4me3 and H3K27me3, respectively (Figure S1C). Additionally, ChIP-sequencing (seq) of Flag-enriched chromatin demonstrated Flag-BAP-H3 incorporation into the genome (Figure 1B and 1D), suggesting that the endogenous N-terminus Flag-BAP tags did not disrupt H3 metabolism.

**Figure 1.**
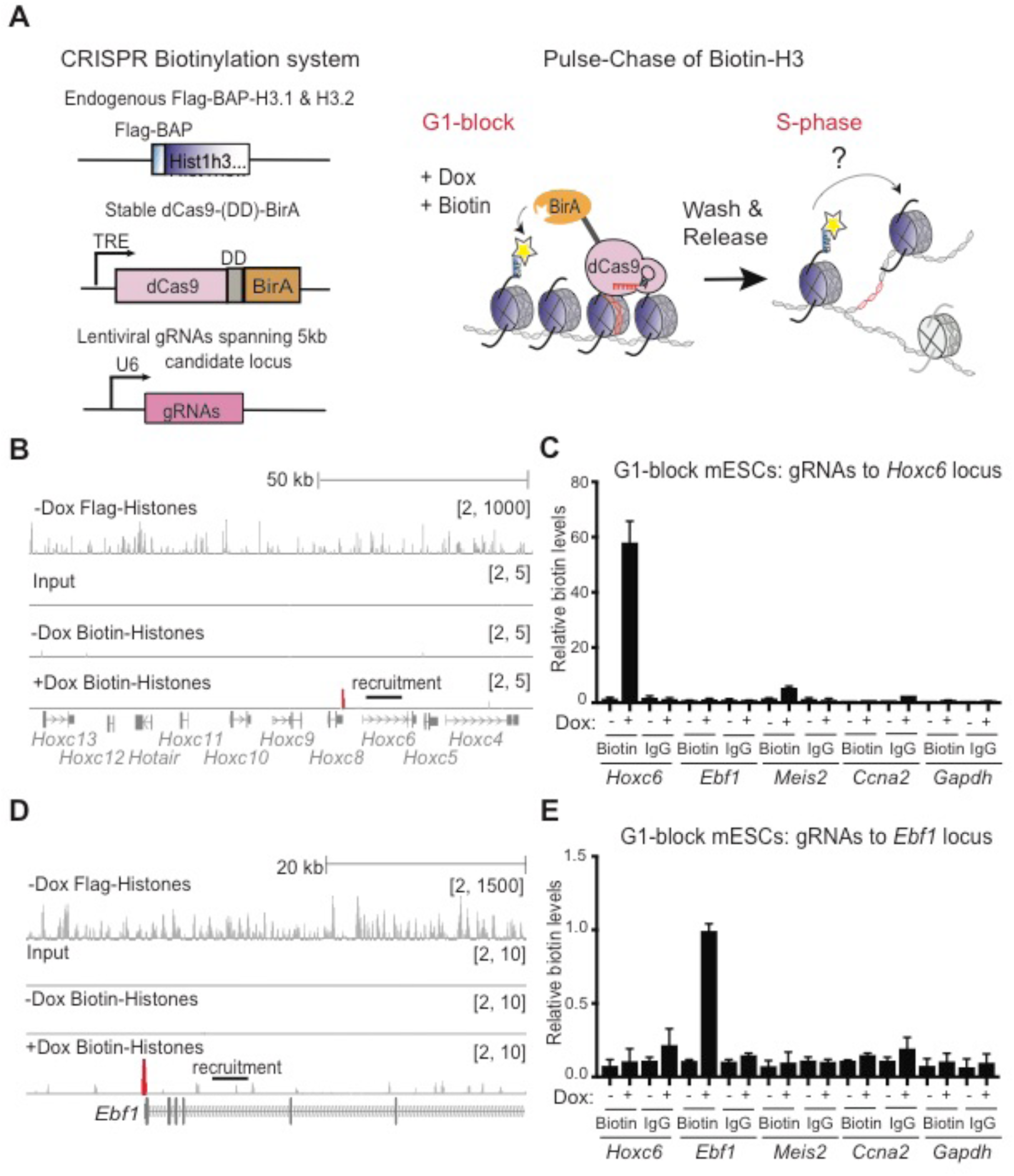
Precise labeling of H3.1 and H3.2 histones in living cells. (**A**) Overview of the system to assess *in vivo* chromatin domain inheritance in mESCs. A master cell line containing endogenous tags of Flag-BAP H3.1 and -H3.2, stable integration of doxycycline (Dox)-inducible dCas9-DD-BirA, and transducible gRNAs spanning 5 kb of a candidate locus is arrested in G1. Following a pulse of doxycycline (Dox) and exogenous biotin, nearby tagged parental nucleosomes are biotinylated (blue histones and yellow asterisks). Wash-off of media releases cells into S-phase wherein the re-distribution of biotin-H3 at a mononucleosomal level is assayed in newly synthesized chromatin. (**B**) Native Flag and biotin ChIP-seq analysis of G1/S-blocked cells at the *HoxC* cluster following dCas9-DD-BirA recruitment. (**C**) Native ChIP-qPCR analysis of biotin-H3 in G1/S-blocked mESCs showing biotin enrichment at the *Hoxc6* locus compared to *Ebf1*, *Meis2*, *Ccna2*, *Gapdh* and IgG controls. (**D**) Flag and biotin native ChIP-seq analysis of cells at the *Ebf1* locus following dCas9-DD-BirA recruitment. (**E**) Native ChIP-qPCR analysis of biotin-H3 in G1/S-blocked mESCs, validating biotin enrichment at the *Ebf1* locus compared to *Hoxc6*, *Meis2*, *Ccna2*, *Gapdh* and IgG controls. Data was normalized to 5% input and error bars represent standard error of three biological replicates.

To spatially recruit dCas9-DD-BirA and biotinylate local parental H3 incorporated into chromatin at a unique desired gene, we stably expressed an array of ~29-35 guide RNAs (gRNAs) spanning 5 kb of sequence from a candidate locus (Figure 1A). We tested the specificity of the system by the introduction of 33 gRNAs in which 5 kb of the *Hoxc6* gene was targeted. We found that during the last step of a double thymidine G1-block synchronization (Figure S1D), a 6 hr pulse with a minimal amount of doxycycline and exogenous biotin followed by a wash-off step was sufficient to observe specific biotinylation of local *Hoxc6* chromatin (Figure 1B and 1C). Briefly, chromatin from G1/S-blocked cells with and without a doxycycline pulse was digested with MNase to obtain mononucleosomes and then biotinylated nucleosomes were isolated by immunoprecipitation using biotin antibodies. Subsequently, native biotin ChIP-seq showed a precise labeling at the *Hoxc6* locus as evidenced by a biotin peak upstream from the 5 kb gRNA recruitment site in doxycycline-treated cells (Figure 1B). Validation of the biotin *Hoxc6* peak through native biotin ChIP-quantitative PCR (ChIP-qPCR) also demonstrated accurate biotinylation of the targeted locus in comparison to nonspecific loci and IgG controls (Figure 1C). Furthermore, correct proximity-based biotinylation in our system was confirmed upon recruitment of dCas9-DD-BirA to a second target region within the *Ebf1* gene (Figure 1D). Native biotin ChIP-seq and ChIP-qPCR again showed a biotin peak upstream from the 5 kb gRNA recruitment site and a specific biotin enrichment of *Ebf1* chromatin in contrast to nonspecific loci and IgG controls (Figure 1D and 1E, respectively). Lastly, to verify the specificity and extent of temporal control on dCas9-DD-BirA expression, we conducted a time course analysis on G1/S-blocked and released ESCs (Figures S2A and 2B, D, F) and observed the targeted recruitment of dCas9 to the *Hoxc6* (and Ebf1 and Meis2, see below) locus and its subsequent dilution after the first cell cycle (Figure S2B). Therefore, our system allows for permanent histone labeling *in vivo* with notable spatial resolution (Figure 1B-E) and tight temporal control of dCas9-DD-BirA recruitment to chromatin (Figure S2B), and results in the precise labeling of parental H3.1 and H3.2 at a single nucleosome level at a desired gene.

### Inheritance of parental nucleosomes from repressive chromatin domains

A rigorous and sustainable repression of key developmental regulators through cell division is required to maintain the ESC pluripotent state (Boyer et al., 2006; Margueron and Reinberg, 2011; Dowen et al., 2014). To determine the localized strategies for the re-deposition of parental nucleosomes following DNA replication, we first assayed the local re-distribution of biotinylated Flag-BAP-H3.1 and -H3.2 at repressed chromatin domains through a timeline of 12, 24, and 48 hr after releasing the ESCs cells from the G1/S-block (Figures 2A and S2A). Among the best studied and developmentally consequential repressed domains in ESCs are *Hox* clusters (Figure S3A) (Boyer et al., 2006). Therefore, we tested parental nucleosome re-deposition by targeting a 5 kb area upstream of the *Hoxc6* gene in dCas9-DD-BirA expressing cells (Figure 1B). These cells were G1/S-blocked and given a 6 hr doxycycline and biotin pulse and subsequent wash-off to label parental nucleosomes. The cells were then released and followed for 12, 24, and 48 hr. Chromatin from these cells was collected, processed through native biotin ChIP, and a 35 kb window spanning 15 kb upstream and downstream from and including the 5 kb recruitment area was assayed for the presence of biotin at a high resolution of 500 bp through qPCR. To quantitatively analyze the parental biotin ChIP-qPCR, we used spike-in *Drosophila melanogaster* chromatin and normalized the data to input, spike-in, and minus-doxycycline native chromatin. Similar to the native biotin ChIP-seq (Figure 1B), *Hoxc6* ChIP-qPCR interrogation of chromatin from cells arrested at the G1/S transition showed a robust biotin peak to the left of the dCas9-DD-BirA recruitment site towards the *Hoxc6* TSS, as well as a less intense peak towards the 3’ end of the transcriptional unit (Figure 2B, 0 hr). This unexpected pattern of nucleosome biotinylation likely depends on the differences in accessibility of BirA to nucleosomes within repressed condensed chromatin, as a different chromatin marking was observed when analyzing euchromatic open chromatin (see below). Nonetheless, the labeling was very specific at the gene level (see Figure 1), allowing us to draw conclusions regarding nucleosome re-deposition at different genes. Subsequent release of the G1/S-blocked ESCs and analysis of the segregation of labeled parental nucleosomes revealed that the biotinylated nucleosomes segregated within the vicinity of the *Hoxc6* locus (Figure 2B, time 12-24 hr) until the signal became undetectable at 48 hr (Figure 2B). Furthermore, quantitative analysis of the highest peak showed a drop in parental biotinylated chromatin enrichment from 1.00 to 0.40 through the first cell division (Figure 2C, Table 1). These findings suggest the positional inheritance of parental nucleosomes to the *Hoxc6* repressed gene in ESCs.

**Table 1:** Quantitative biotinylated nucleosome remaining associated with chromatin across DNA duplication

| Gene Name | Chromatin state | % Biotin Remaining |  |
| --- | --- | --- | --- |
| | | $\Delta t$ 0-12 hr | $\Delta t$ 0-24 hr |
| *Hoxc6* | Repressed | 40% | 31% |
| *Ebfl* | Repressed | 50% | 39% |
| *Meis2* | Repressed | 63% | 32% |
| *Gata2* (-RA) | Repressed | 34% | 10% |
| *Gata6* (-RA) | Repressed | 57% | 20% |
| *Ccna2* | Active | 12% | 4% |
| *Pou5f1* | Active | 7% | 1% |
| *Nanog* | Active | 20% | 1% |
| *Gata2* (+RA)* | Active | 0% | 0% |
| *Gata6* (+RA)* | Active | 8% | 1% |

\*
Candidate loci analyzed (left column) for the percentage of biotin retention 12 and 24 hr after released from the G1 block and within S-phase of the cell-cycle (right columns). The active and repressed genes analyzed are indicated (middle column). The *Gata* loci in which Retinoid Acid (RA) was added to induced their expression is indicated. The expression of the other genes analyzed was analyzed at steady-state conditions.

**Figure 2.**
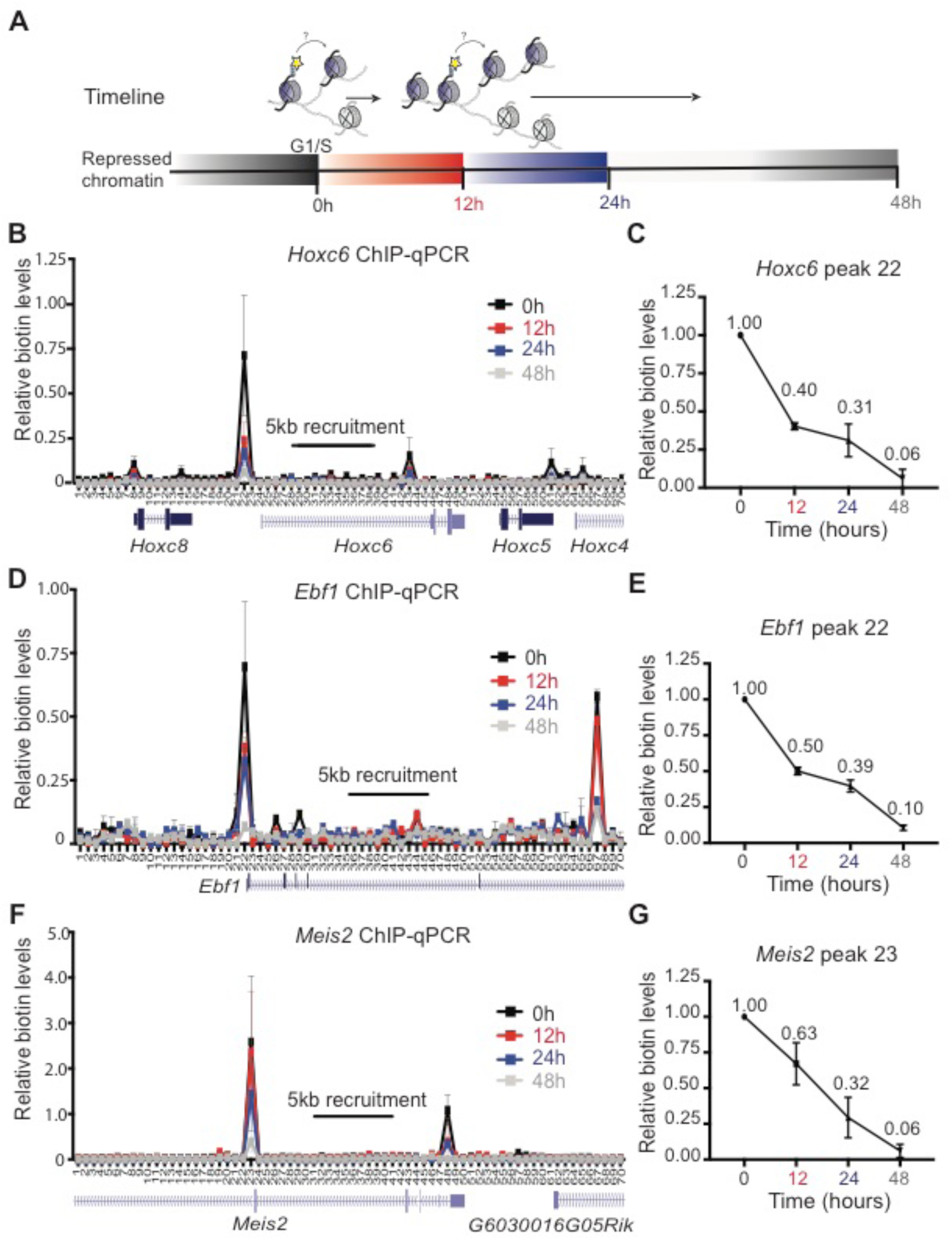
Repressed parental H3 domains are re-distributed locally. (**A**) Experimental timeline whereby ESCs are synchronized at G1/S phase (0 hr) and released to continue the cell cycle. Cells were harvested at 12 hr, 24 hr, and 48 hr intervals thereafter. (**B-G**) Native mononucleosomal biotin ChIP-qPCR in G1/S-blocked and released mESCs following a 6 hr pulse of Dox and exogenous biotin in cells targeting BirA to the *Hoxc6* (**B, C**), *Ebf1* (**D, E**), and *Meis2* (**F, G**) loci. Data shows the average of 3 biological replicates spanning a 35 kb area at a resolution of 500 bp. (**C**), (**E**), and (**G**), Graph highlighting the highest peak of corresponding assays: *Hoxc6* primer set 22 (**B**), *Ebf1* primer set 22 (**D**), and *Meis2* primer set 23 (**F**). All biotin enrichment levels are relative to input, normalized to *Drosophila* chromatin spike-in, followed by subtraction of the minus-Dox (-Dox) control. Error bars represent standard error. For panels C, E, and G, the dataset for 0 hr was set to 1.

To further analyze whether the observed positional inheritance of the *Hoxc6* repressed domain was a general phenomenon across repressed chromatin domains, dCas9-DD-BirA was similarly recruited to two other transcriptionally inactive domains on separate chromosomes (Figure S3B and S3C). We selected gRNAs to target a 5 kb region in the *Ebf1* or the *Meis2* gene and observed an analogous biotin enrichment at the 5’ and 3’ recruitment areas in G1/S-blocked ESCs (Figure 2D and 2F, respectively, 0 hr). Similar to the *Hoxc6* locus, DNA replication resulted in the dilution of the parental biotin signals within their respective regions (Figure 2D and 2F, 12-48 hr) and quantitative analysis of the most intense peak at each locus showed a drop in parental biotinylated chromatin enrichment from 1.00 to either 0.50 or 0.63 through the first cell division (Figure 2E and 2G, Table 1). These observations from three independent repressed chromatin domains point to parental histones being re-deposited locally.

### Lack of inheritance of parental nucleosomes from active chromatin domains

That parental histones were retained locally during replication of repressed chromatin domains warranted the comparable analysis of active chromatin domains. We employed gRNAs tiling to a 5 kb region upstream of the *Ccna2* gene and during the final step of double thymidine synchronization, induced the transient expression of dCas9-DD-BirA and subsequent biotinylation of Flag-BAP-H3.1 and -H3.2 at *Ccna2* chromatin, as performed above. We then assayed the local re-distribution of biotinylated Flag-BAP-H3.1- and -H3.2-labeled nucleosomes and the results showed a lower level of nucleosome labeling and a much broader area from the *Ccna2* 5 kb recruitment site (Figure 3A, 0 hr), likely reflecting that the chromatin is “open” and more accessible to the dCas9-DD-BirA. Subsequent to the release from G1/S-blocked *Ccna2*, the biotin enriched parental histones could not be detected after DNA replication (Figure 3A, 12 hr). Quantitative analysis of the highest biotin peak surrounding the *Ccna2* 5 kb recruitment area showed a drastic drop in parental biotinylated chromatin enrichment from 1.00 to 0.12 through the first cell division (Figure 3B, 12 hr and Table 1). These data point to the non-local re-distribution of parental histones in the *Ccna2* active domain.

**Figure 3.**
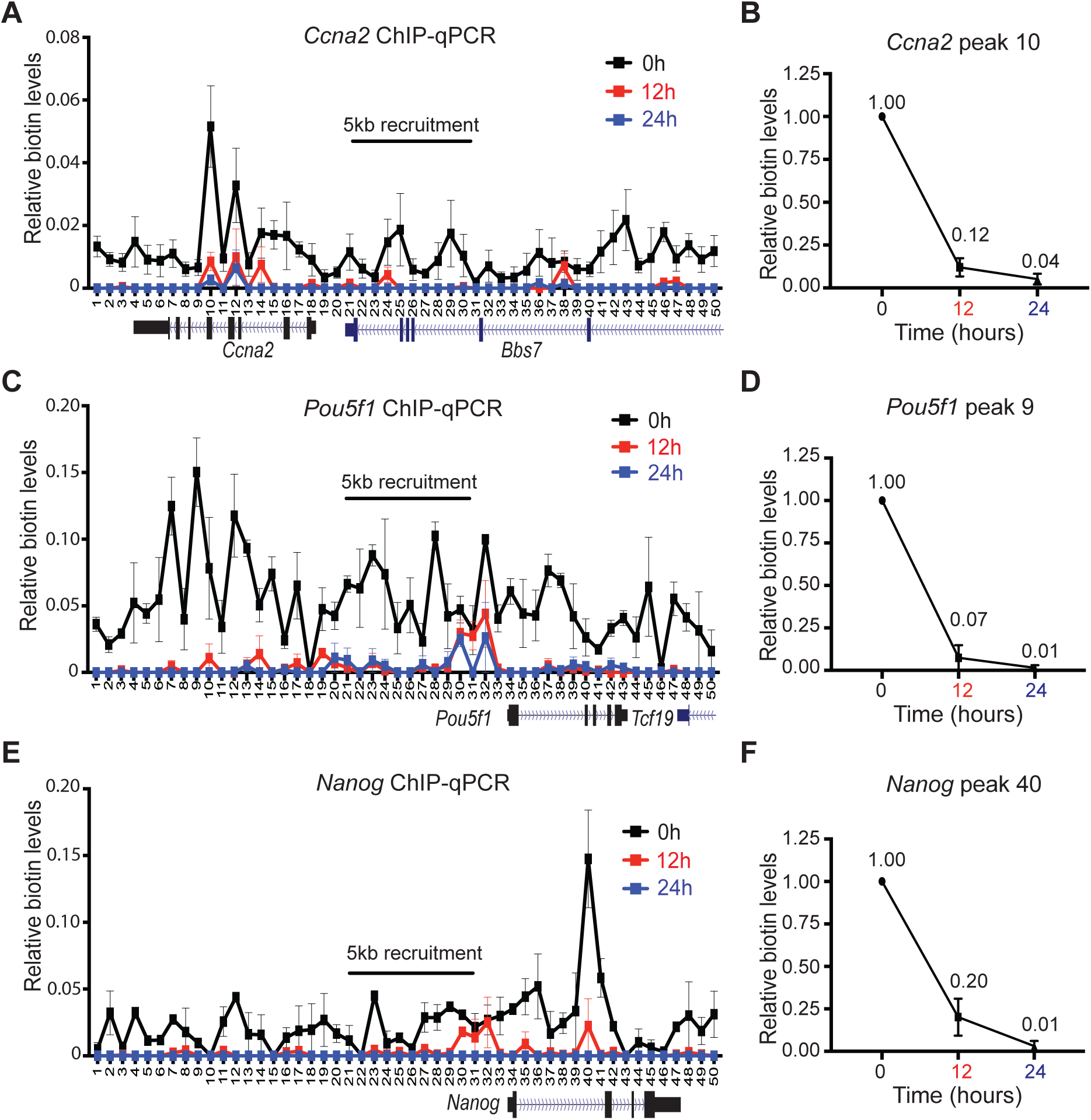
Dynamic distribution of active parental H3 domains. Native mononucleosomal biotin ChIP-qPCR from G1/S-blocked and released mESCs following a 6 hr Dox and exogenous biotin pulse in cells targeting BirA to the *Ccna2* (**A, B**), *Pou5f1* (**C, D**), and *Nanog* (**E, F**) loci. Data shown is the average of 3 biological replicates spanning a 25 kb area at a resolution of 500 bp. (**B**) (**D**), and (**F**), graph is highlighting the highest peak of corresponding assays: *Ccna2* primer set 10 (**A**), *Pou5f1* primer set 9 (**C**), and *Nanog* primer set 40 (**E**). All biotin enrichment levels are relative to input, normalized to *Drosophila* chromatin spike-in, followed by subtraction of the -Dox control. Error bars represent standard error. For panels **B**, **D**, and **F**, the dataset for 0 hr was set to 1.

As active domains replicate earlier in S-phase (Rhind and Gilbert, 2013), we examined whether the loss of *Ccna2* biotinylated chromatin occurs as early as 6 hr, a time at which mESCs are in mid S-phase prior to cell division (Figure S4A). Indeed, native biotin ChIP-qPCR of *Ccna2* chromatin showed a drastic decrease in the biotin signal at this time point and a more pronounced loss at 12 hr (Figure S4B, 6 hr and 12 hr). Furthermore, transcription inhibition during S-phase progression was ineffectual with respect to the decrease in the biotin signal (Figure S4B, 6hr + Triptolide), suggesting that the loss in parental H3 in *Ccna2* active domain is not due to transcriptional turnover. Thus, biotinylated parental histones, at least within the *Ccna2* locus, distribute non-locally upon DNA replication, revealing an unexpected, but biologically sound difference in the spatial inheritance of nucleosomes at active versus repressed loci (see Discussion), which could not be appreciated with previously published genome-wide approaches.

To interrogate whether this non-local distribution of parental histones is specific to the *Ccna2* locus or represents a wider phenomenon of nucleosome inheritance associated with active domains, we expanded our studies and analyzed the loci of pluripotent factors *Pou5f1* and *Nanog*, which are highly expressed in ESCs and encode transcription factors critical for ESC pluripotency and self-renewal capacity (Takahashi and Yamanaka, 2006). Targeting dCas9-DD-BirA to either of these loci in G1/S-blocked cells again resulted in a broader biotin enrichment surrounding the 5 kb gRNA recruitment area (Figure 3C and 3E, respectively). Furthermore, and similar to the case of chromatin associated with the *Ccna2* locus, the release of G1/S-blocked biotinylated *Pou5f1* or *Nanog* chromatin led to a loss in biotin signal upon the first cell division (Figure 3C and 3E, respectively). We quantitatively assessed the reduction of the biotin signal from the peaks of *Pou5f1* and *Nanog*, which again revealed a drastic drop in biotin enrichment from 1.00 to 0.07 for *Pou5f1* (Figure 3D, 12 hr) and from 1.00 to 0.20 for *Nanog* (Figure 3F, 12 hr) within the first cell division (Table 1). The loss in biotin signal following DNA replication in active chromatin domains suggests that parental H3.1- and H3.2-containing nucleosomes are randomly re-deposited within active genes. Alternatively, the loss in biotin signal could be due to a trivial loss in detection. The latter is not likely given the results obtained with two other active domains, as shown below. These results point to the distinction between nucleosome re-deposition in active versus repressed chromatin, with biotinylated parental histones being dispersed in the case of active domains and positionally inherited in the case of repressed ones.

### The segregation of parental nucleosomes requires DNA replication

To assess whether DNA replication is required for the local and non-local segregation of parental nucleosomes associated with repressed and active chromatin domains, respectively, we examined biotin signal dilution when DNA replication is blocked. To this end, shortly after cells were released from the G1/S-blocked, biotinylated nucleosomes present on the *Hoxc6* and *Pou5f1* cells were kept in S-phase within a 12 or 24 hr timeline of Aphidicolin treatment (Figures 4A and S5). The peaks at the repressed *Hoxc6* and active *Pou5f1* loci enriched for biotin (Figures 2C and 3D), significantly sustained biotin enrichment in the presence of Aphidicolin (Figure 4B-E). Indeed, the absence of biotin dilution in Aphidicolin-treated *Hoxc6* and *Pou5f1* chromatin suggests that the local dynamics of parental nucleosome segregation were frozen when S-phase progression was blocked.

**Figure 4.**
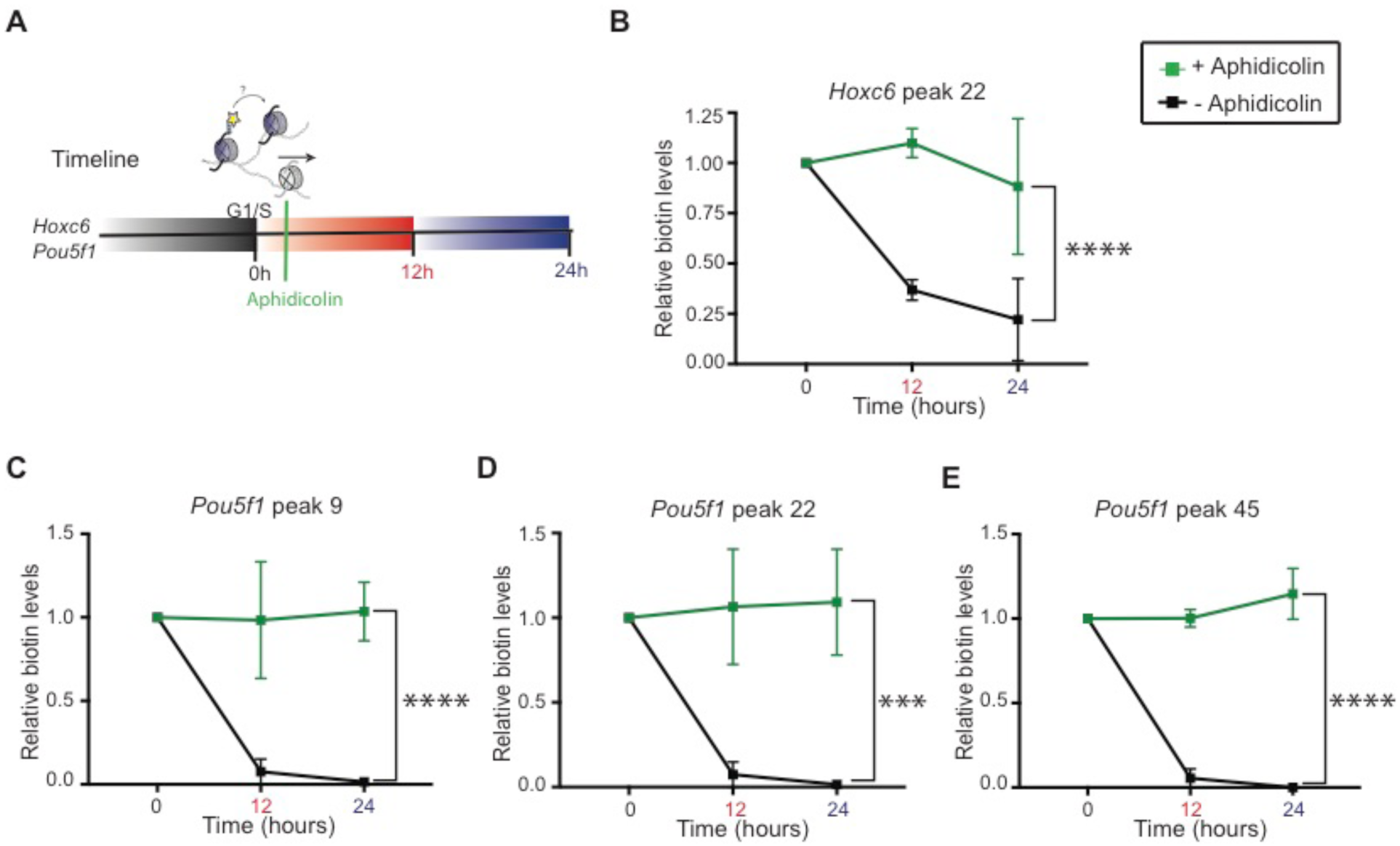
Parental H3 nucleosome segregation is replication dependent. (**A**) Experimental timeline whereby ESCs are synchronized at G1/S phase (0 hr) and released to continue the cell cycle or treated with 1 mg/ml Aphidicolin to block S-phase progression. Cells were harvested at 12 hr and 24 hr thereafter. (**B-E**) Native biotin ChIP-qPCR analysis of the *Hoxc6* peak primer set 22 (**B**), *Pou5f1* peak primer set 9 (**C**), *Pou5f1* peak primer set 22 (**D**), and *Pou5f1* peak primer set 45 (**E**) in cells treated (green) or not treated (black) with Aphidicolin following biotin enrichment during G1 synchronization and release into S-phase. All biotin enrichment levels are relative to input and normalized to *Drosophila* chromatin spike-in followed by subtraction of the minus-Dox (-Dox) control. Biotin-enriched Aphidicolin chromatin was then normalized to biotin-H3 at 0 hr. Error bars represent standard error of three biological replicates. Statistical significance determined using 2way ANOVA (*** p<0.001 and **** p<0.0001).

### Inheritance of repressed chromatin domains as a function of cellular differentiation

To maintain a pluripotent state, ESCs require that genes encoding lineage-specifying developmental regulators remain repressed (Figure S6), as their specific expression will promote differentiation to a specific lineage and consequently a change in cellular identity (Boyer et al., 2006; Margueron et al., 2009; Dowen et al., 2014). The GATA factors are such genes whose expression in ESCs leads to the differentiation of mesoderm and ectoderm-derived tissues (Lentjes et al., 2016). Since the retinoic acid (RA) signaling molecule can activate the expression of GATA factors (Zhang et al., 2015), we analyzed the segregation of parentally marked-nucleosomes upon RA treatment. As expected, RA addition to ESCs for 12 hr prior to G1-synchronization and release (Figure 5A), resulted in the induction of *Gata2* and *Gata6* gene expression (Figure 5B). Introducing gRNAs to target a 5-kb area upstream of the *Gata2* and *Gata6* genes resulted in the recruitment of dCas9-DD-BirA and the subsequent biotinylation of parental H3.1 and H3.2-containing nucleosomes (Figures 5C and D). As expected from our results with repressed chromatin domains (Figure 2), we found a discrete peak in the *Gata2* and *Gata6* loci of G1/S-blocked chromatin in cells not treated with RA (Figure 5C and D, 0 hr, -RA). Similarly, the subsequent release of these G1/S-blocked ESCs showed the re-deposition of biotin parental nucleosomes within the vicinity of the initial *Gata2* and *Gata6* peak position (Figure 5C and D, 12-24 hr in -RA).

**Figure 5.**
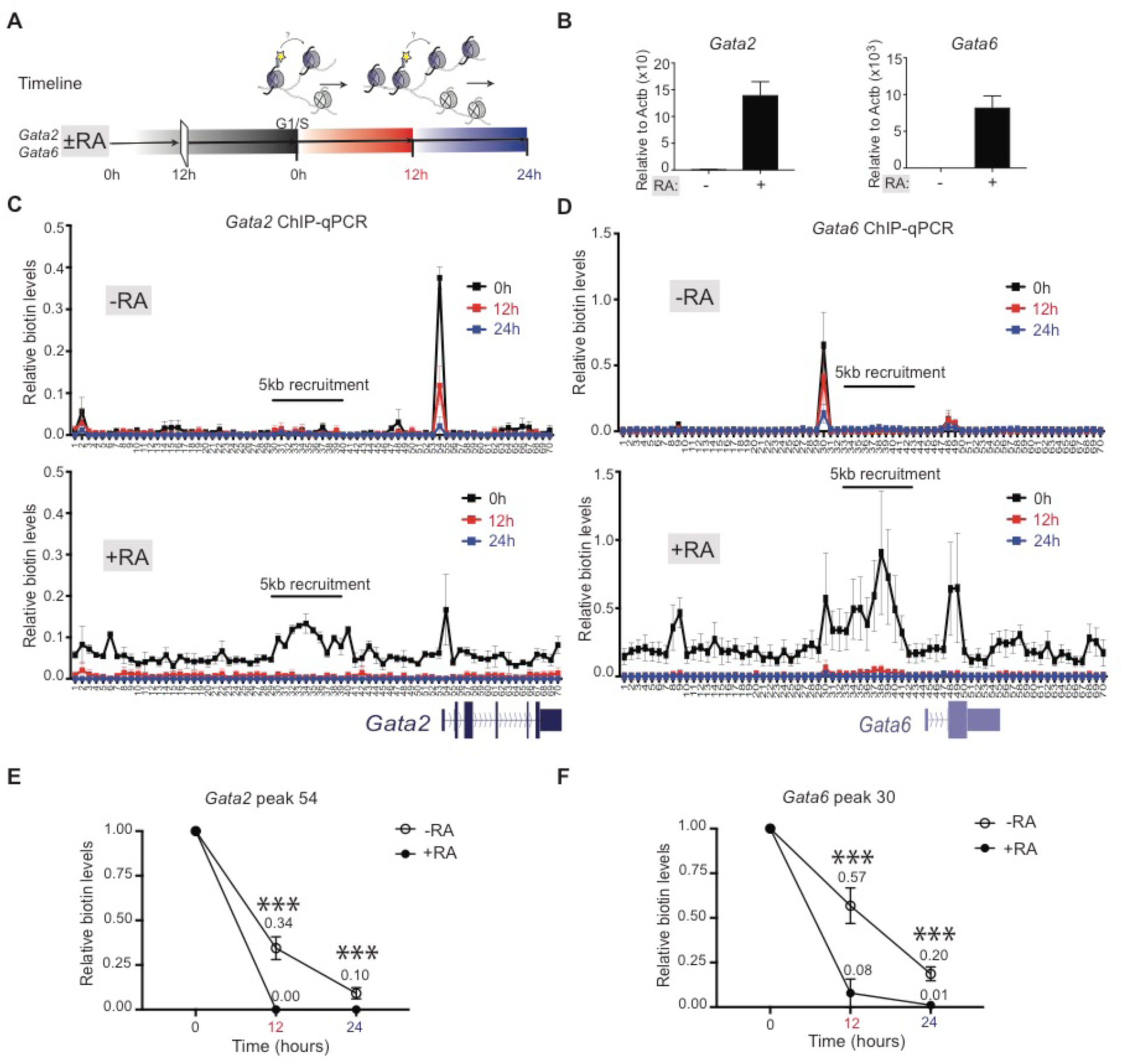
Altering H3 repressed to active domains changes local parental H3 recycling. (**A**) Experimental timeline whereby ESCs are treated or not treated with RA for 12 hr prior to synchronizing at G1/S phase (0 hr) and released to continue the cell cycle. Cells were harvested at 12 hr and 24 hr thereafter. (**B**) Relative mRNA expression of *Gata2* and *Gata6* genes in ESCs treated or not treated with RA for 48 hr. Data was normalized to Actin expression and minus RA control. (**C-D**) Native mononucleosomal biotin ChIP-qPCR in G1/S-blocked and released mESCs treated or not treated with RA following a 6 hr pulse of Dox and exogenous biotin in cells targeting BirA to the *Gata2* (**C**) and *Gata6* (**D**) loci. Data shows the average of 3 biological replicates spanning a 35 kb area at a resolution of 500 bp. (**E-F**) Graph highlighting the highest peak of corresponding assays: *Gata2* primer set 54 (**E**) and *Gata6* primer set 30 (**F**). Biotin enrichment levels are relative to input and normalized to *Drosophila* chromatin spike-in followed by subtraction of the minus-Dox (-Dox) control, then normalized to biotin-H3 at 0 hr. Error bars represent standard error of three biological replicates. Statistical significance determined using 2way ANOVA (_***_ p<0.001).

Upon RA-induced transcription activation of *Gata2* and *Gata6* (Figure 5B), the levels of biotinylated parental H3.1 and H3.2 chromatin remained comparable to that of a repressed state, but the biotin enrichment across *Gata2* or *Gata6* was now broader (Figure 5C and D, 0 hr in +RA). Of note, steady-state active chromatin (*i.e. Ccna2*, *Pou5f1* and *Nanog*) exhibited a lower and much broader biotin signal (Figure 3) than observed when *Gata2* and *Gata6* loci were captured immediately upon RA-induced activation. Yet, similar to the non-local segregation of histones in steady-state active chromatin (Figure 3), *Gata2-*and *Gata6-*expressing cells showed a loss in biotinylated parental H3 chromatin at these loci (Figure 5C and D, 12-24 hr in +RA). Quantitative assessment for the distribution of the original biotin peak observed in repressed chromatin shows that biotinylated parental nucleosomes were diluted from 1.00 to 0.34 for *Gata2* (Figure 5E, 12 hr, -RA) and from 1.00 to 0.57 for *Gata6* (Figure 5F, 12 hr, -RA). In comparison, a drastic drop in biotin signal was observed for active chromatin as biotin-labeled nucleosomes changed from 1.00 to 0.00 for *Gata2* (Figure 5E, 12 hr, +RA) and from 1.00 to 0.08 for *Gata6* (Figure 5F, 12 hr, +RA, Table 1). These results argue for the existence of distinct localized strategies for the re-deposition of parental nucleosomes in repressed versus active chromatin as changing the transcriptional state of *Gata2* and *Gata6* genes significantly altered the dynamics of parental H3 segregation (Figure 5E and 5F, ±RA).

## Discussion

A key feature to the control of gene expression is embodied in the histones and their associated PTMs that modulate the structure, stability, and dynamics of the local chromatin state (Luger et al., 2012; Stillman, 2018). Although it is accepted that histone PTMs selectively impact a transcriptional program, it is unclear if histones and their PTMs are determinants to the heritable transmission of an ‘epigenetic’ state from one cell generation to the next. To address this question, we developed a system in which pre-existing H3.1- and H3.2-bearing nucleosomes are marked at a precise single gene and followed across DNA replication to assess their local segregation in active or repressed chromatin domains. We find a spatial conservation in the recycling of intact parental H3.1- and H3.2-nucleosomes within repressive chromatin domains, but this local re-deposition is absent in the case of nucleosomes associated with active chromatin (Figure 6 and Table 1). In the latter case, the chromatin is open (euchromatin), as evidenced by the broader area and decreased signals in our nucleosome labeling experiments of steady-state active genes (*i.e. Ccna2*, *Pou5f1* and *Nanog*). In contrast to repressed chromatin (*e.g. Hoxc6*), this biotin-H3 enrichment is lost upon DNA replication (Figure 3, Table 1). When biotin enrichment levels of newly activated chromatin (*i.e. Gata2* and *Gata6* loci) are comparable to those of their respective, repressed state (Figure 5D and 5E, ±RA 0 hr), a loss in biotin-H3 in active chromatin is again observed, in contrast to their repressed states that retain the parental histone signals (Figure 5D and 5E, ±RA 12 hr, Table 1). Thus, re-deposition of parental H3.1- and H3.2-nucleosomes in active chromatin domains lack heritable transmission. We conclude that nucleosomes in a repressive chromatin state segregate locally such that this signature is conveyed directly to the same domains on daughter DNA. In the case of active domains, we propose that the associated histone PTMs function solely to facilitate transcription and can be readily restored subsequent to DNA replication upon re-engagement of the appropriate sequence-specific transcription factors to direct transcription (Ptashne and Gann, 2002; Reinberg and Vales, 2018).

**Figure 6.**
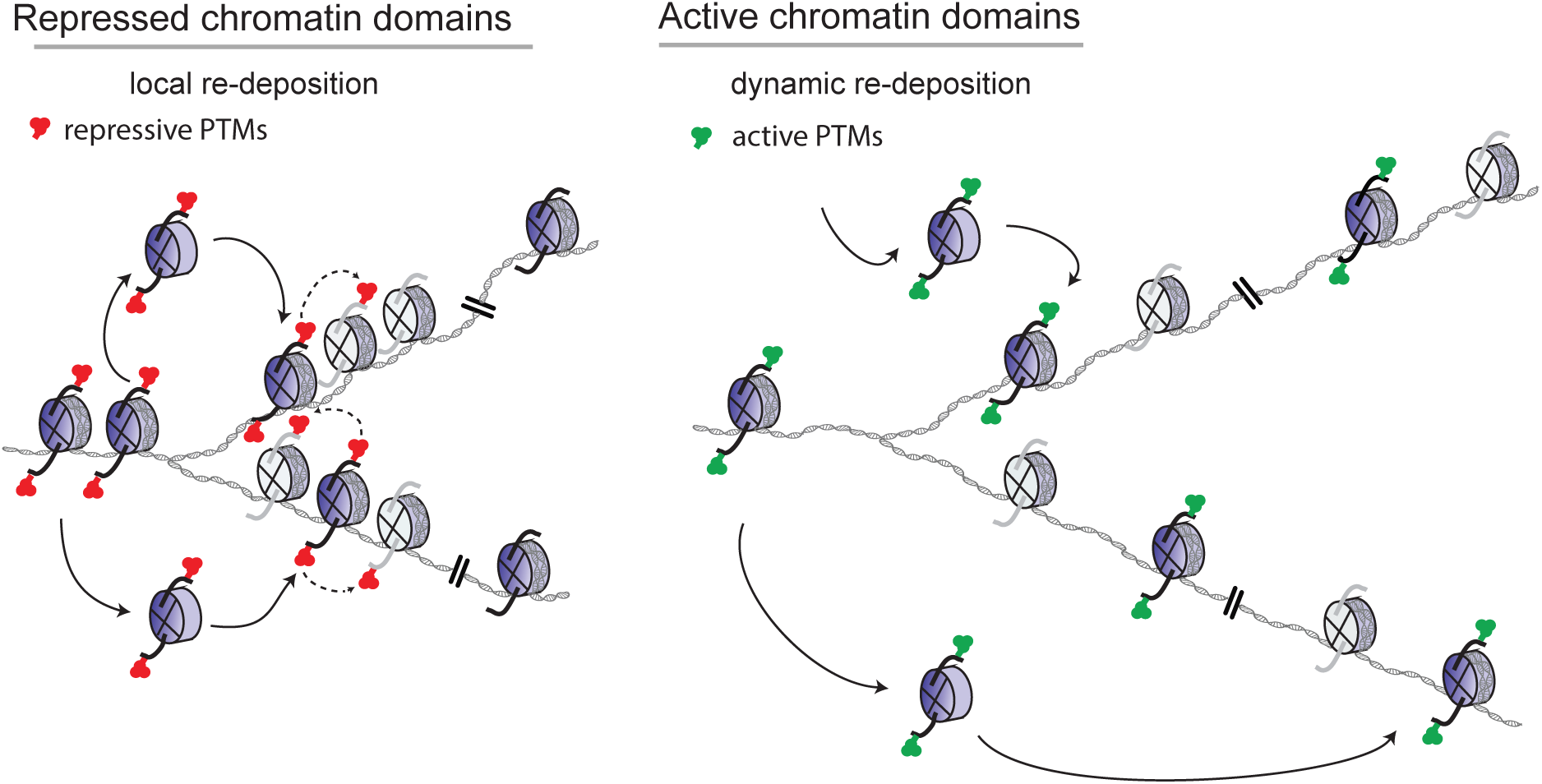
Parental H3 nucleosomes segregate locally in repressed chromatin domains. In repressed chromatin domains, a degree of spatial conservation in the re-deposition of intact pre-existing (H3.1-H4)_2_ or (H3.2-H4)_2_ tetrameric core-bearing histone PTMs (blue histones and solid arrows) are sufficient to transmit chromatin states to daughter cells if maintenance enzymes are available and can restore the transmitted modification(s) to neighboring newly synthesized histones (gray histones and dashed arrows) (Reinberg and Vales, 2018). This local re-deposition of parental H3.1 or H3.2 nucleosomes is dynamic in active chromatin domains (see Discussion).

In a prokaryotic cell, gene regulation is not conveyed by structural changes to the DNA and its histone-like proteins (Ptashne and Gann, 2002; Dillon and Dorman, 2010). Instead, the genome is open and subject to the function of activators and repressors that control inducible operators to alter gene transcription (Ptashne and Gann, 1997; Ptashne and Gann, 2002). It is not until the advent of unicellular eukaryotes that histones and in particular, their amino-terminal tails afford a link to the direct regulation of gene transcription (Stillman, 2018). A key example is histone H4 and its acetylation at lysine 16 (H4K16ac), which facilitates open higher order chromatin structures which is conductive to gene expression (Shogren-Knaak et al., 2006). However, similar to prokaryotes, unicellular eukaryotes require activators to initiate gene transcription. In multicellular organisms, master regulators establish transcriptional outputs through their sequence-specific DNA binding activity, which recruits coactivators and RNA polymerase II to promoter sequences (Patikoglou and Burley, 1997). RNA Polymerase II in turn recruits independent methyltransferases that methylate histone H3 at lysine 4 and 36. Methylated H3K4 and H3K36 are associated with active chromatin, functioning in combination with other proteins that recognize the methyl groups and further facilitate transcription (Sims et al., 2004). Thus, in eukaryotes, histones and their associated PTMs evolved to enable transcriptional changes, but sequence-specific transcription factors are required for transcription as in the case of prokaryotic cells and the histone modifications associated with active chromatin are a consequence of transcription activity.

Strikingly, it is through gene repression that histone PTMs show a heritable gene regulatory system in eukaryotic cells. For example, in *S. cerevisiae*, the repression of the silent mating type loci *HMLα* and *HMRa* requires the deacetylation of histone H4K16ac that is catalyzed by the silent information regulator (Sir) complex (Sir2-Sir3-Sir4), initially recruited through its interaction with the Sir1 adaptor (Johnson et al., 1990; Stillman, 2018). While Sir3 and Sir4 foster the expansion of this silencing, the histone deacetylase activity of Sir2 maintains the inheritance of the deacetylated version of H4K16 through multiple cell divisions in the absence of the initial signal from Sir1 (Johnson et al., 1990; Allshire and Madhani, 2018; Stillman, 2018).

In multicellular organisms, histone PTMs associated with repressed chromatin also mediate a heritable chromatin state. These heritable gene silencing histone PTMs include H3K9me3 catalyzed by Suv39h1/h2 and H3K27me2/me3 catalyzed by the Polycomb Repressive Complex 2 (PRC2). Both Suv39h1/h2 and PRC2 exhibit a *bone fide* “write and read” mechanism by which the polypeptide or complex, respectively, deposits their corresponding PTM and binds to the PTM resulting in an allosteric stimulation of the enzymatic activity (Margueron et al., 2009; Allshire and Madhani, 2018; Oksuz et al., 2018; Reinberg and Vales, 2018). Similar to Sir2 in yeast, both SUV39H1 and PRC2 can propagate and maintain a repressive chromatin state through multiple cell divisions in the absence of the initiating signal; an epigenetic phenomenon (Allshire and Madhani, 2018; Reinberg and Vales, 2018).

In addition to this “write and read” mechanism, the inheritance across DNA replication of a repressed chromatin state entails that the histone methyltransferase activity be in proximity to parental nucleosomes such that their associated PTMs are transmitted as newly synthesized histones (naïve nucleosomes) comprise half of replicative chromatin and might attenuate a repressed state. Most importantly, the pre-existing histones and their PTMs must segregate to the same domains on daughter DNA strands. Therefore, our aim was to assess the segregation dynamics of parental H3.1 and H3.2-containing nucleosomes by establishing a system that precisely and irreversibly labels nucleosomes at a specific location and at a specific time within a single gene. Our system allows for parental H3.1 and H3.2-nucleosomes to be biotinylated prior to DNA replication and its tagged density to be followed through chromatin duplication. It is only through our technique that accurately scores for parental nucleosome position prior to DNA replication that the parental histone origin can be determined. Although ChIP-seq and imaging systems in conjunction with synchronization experiments can detect the restoration dynamics of newly synthesized chromatin (Clement et al., 2018; Reveron-Gomez et al., 2018), such genome-wide approaches cannot ascertain the original locale of a parental nucleosome. Thus, our findings here can attribute the inheritance of a repressed state to both the local segregation of parental nucleosomes and to the “write and read” modules of the histone methyltransferase that deposit the repressive modification (Figure 6).

A feasible explanation for this local conservation of repressed chromatin domains versus active chromatin domains is that replication of repressed chromatin occurs late in S phase, in contrast to that of active chromatin, which is an early event. This timing difference is known to affect the rate of replication, such that euchromatin is replicated at a faster rate than heterochromatin (Rhind and Gilbert, 2013). The slower rate in the latter case, might facilitate parental histone segregation to the same domains. Whether these differences can account for the observed positional inheritance of repressive chromatin domains remains to be elucidated, but as replication timing and chromosomal domain structures are intertwined (Rhind and Gilbert, 2013), it is possible that active and repressed chromatin form self-interacting domains that set thresholds for the accessibility of factors promoting the appropriate chromatin organization during DNA replication. We speculate that distinct chaperones might operate during late S-phase, but are absent in early replicating chromatin. Our findings set the stage for investigating the veracity of such scenarios as they clearly demonstrate a fundamental feature of epigenetic inheritance: that nucleosomes associated with repressed chromatin segregate to the same chromatin domains whereas those associated with active chromatin exhibit a dispersed re-distribution.

## Acknowledgments

We thank Drs. S. Krishnan, E. Campos, and S. Tu for thoughtful discussions. We thank Dr. L. Vales for discussion and revision of the manuscript and D. Hernandez for excellent technical assistance. We also are grateful to Brian J. Abraham and Ann Boija for ChIP-seq discussions. We are also appreciative for the expertise offered at the following facilities at New York University Langone Health: the Genome Technology Center for help with sequencing and the Cytometry and Cell Sorting core facility for cell sorting/flow cytometry technologies.

## Funding

This work in D.R. Lab was supported by the Howard Hughes Medical Institute and the National Cancer Institute (NCI NIH 9R01CA199652-13A1). T.M.E was supported by NCI NIH 3R01CA199652-14S1. The NYU Langone’s Cytometry and Cell Sorting Laboratory is supported in part by grant P30CA016087 from the National Institutes of Health/National Cancer Institute.

## Author contributions

T.M.E., R.B. and D.R. conceptualized the project. T.M.E. performed the experiments and wrote the paper. O.O. helped with formal analysis and development of methodology. N.D. performed bioinformatic analysis. D.R. supervised the study.

## Competing interests

Danny Reinberg is a cofounder of Constellation Biotechnology and Fulcrum Biotechnology

## STAR METHODS

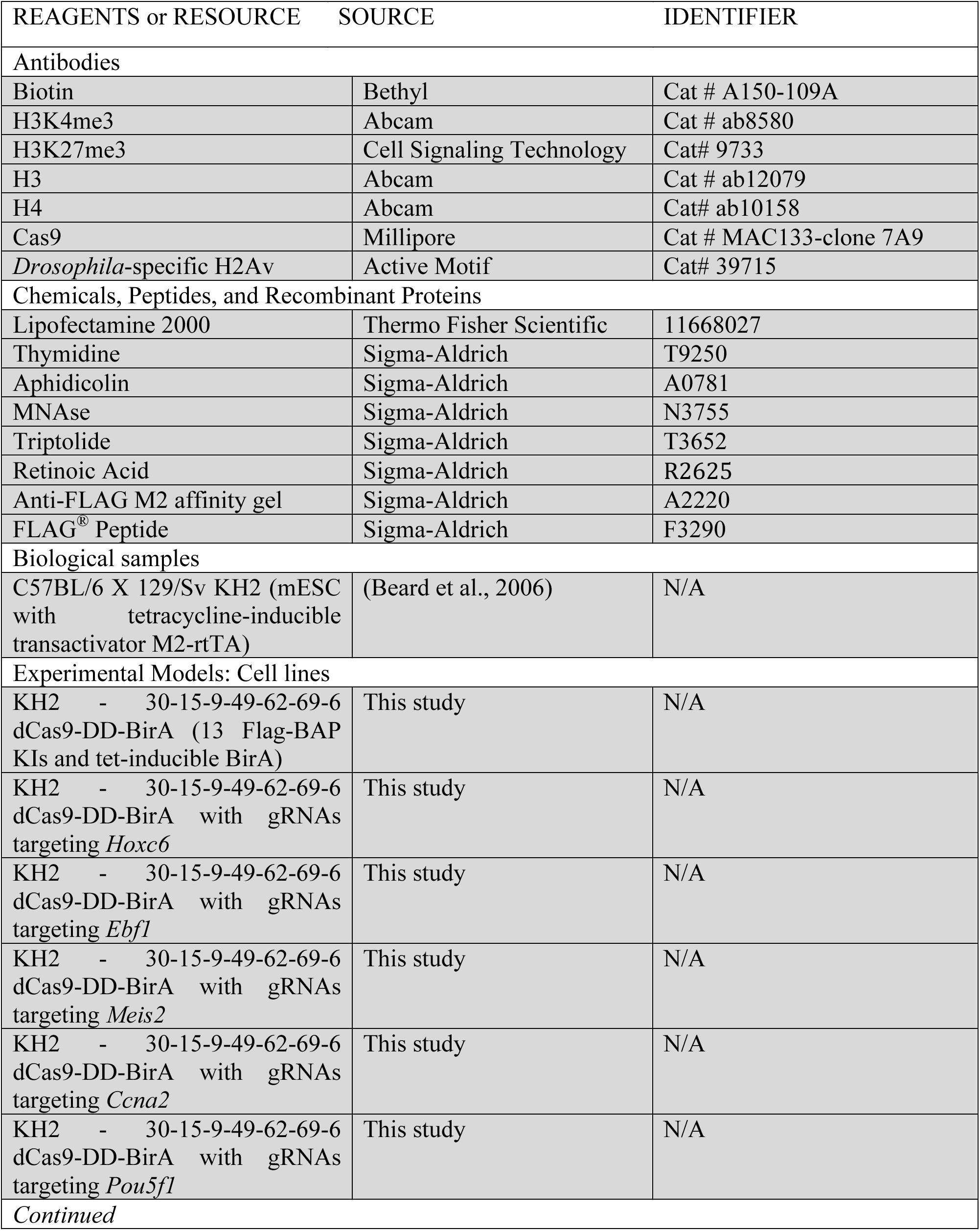
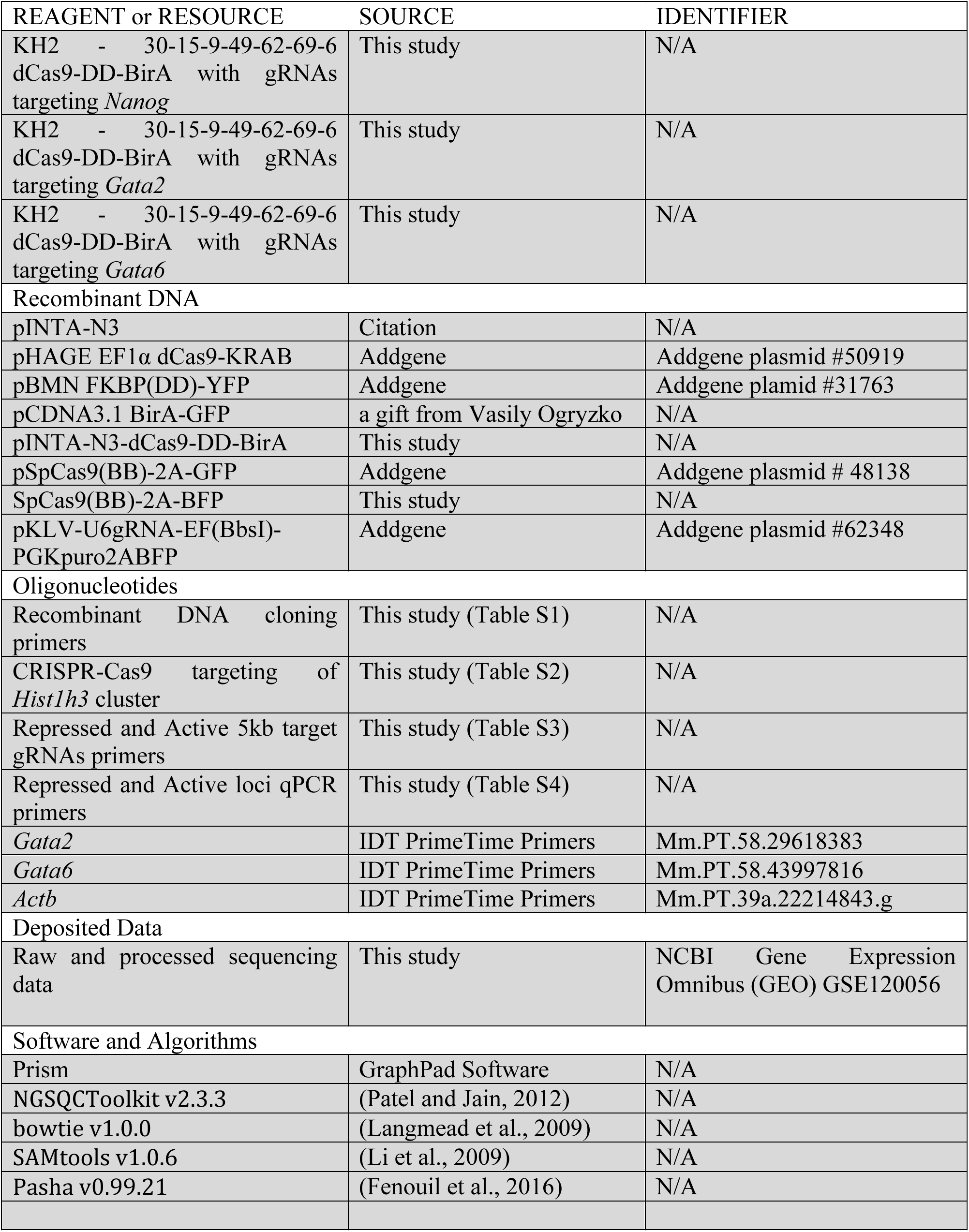

## CONTACT FOR REAGENT AND RESOURCE SHARING

Further information and requests for resources and reagents may be directed to the Lead Contact, Danny Reinberg.

## EXPERIMENTAL MODEL AND SUBJECT DETAILS

The following were used in this study: KH2 mouse ESCs and Human embryonic kidney 293T (293T) cells. KH2 ESCs were grown in DMEM supplemented with 15% FBS (for CRISPR/Cas9 targeting) or Tet-free FBS (Gemini 100-800, for biotinylation experiments), L-glutamine, penicillin/streptomycin, non-essential amino acids, 0.1 mM β-mercaptoethanol, LIF, and 2i inhibitors which include 1 µM MEK1/2 inhibitor (PD0325901) and 3 µM GSK3 inhibitor (CHIR99021) on 0.1% gelatin coated plates. 293T cells were grown in DMEM supplemented with 10% FBS, L-glutamine and penicillin/streptomycin.

## METHOD DETAIL

### Cell Line Generation

To generate dCas9-DD-BirA expressing stable cell lines, 2 µg of pINTA-dCas9-DD-BirA plasmid was transfected with Lipofectamine 2000 (Thermo Fisher Scientific) into KH2 ESCs. After 2 weeks of 200 µg/ml of Zeocin selection, single colonies were selected. dCas9-DD-BirA inducibility was tested and cells were provided with FUCCI reporter (Sladitschek and Neveu, 2015) for testing in synchronization experiments. The desired stable clone was expanded for CRISPR/Cas9 targeting of Flag-BAP knockins to H3.1 and H3.2 copies in the *Hist1h3* cluster (see below).

### Cell Synchronization and *in vivo* biotinylation

For G1 synchronization experiments, ESCs were plated at 50-60% confluency and pulsed for 18 hr with 8 µM thymidine, followed by a PBS wash and a 7-8 hr release into 2i Tet-free media. mESCs were then given a second thymidine treatment at a 5-6 µM concentration for 12-13 hr. The G1-block was confirmed by staining DNA with propidium iodide. For the biotinylation experiments, a 6 hr pulse of 2 µg/ml of doxycycline and 1 µg/ml of exogenous biotin was given during the latter half of the second thymidine treatment. Next, ESCs were either harvested at the G1-block (starting time point, 0 hr) or washed with PBS and released to 2i Tet-free media for the indicated time points. In S phase-block experiments, 1 µg/ml of Aphidicolin was added to the 2i Tet-free media following release from the G1-block and for the transcription inhibition experiments, 1 mM of Triptolide was added following release into S-phase. Lastly, for Retinoic Acid (RA) differentiation, mESCs were given 1 µM RA 12 hr prior to double thymidine synchronization and kept in RA-media throughout G1-synchronization and experimental time course.

### Cloning

Cloning of the pINTA-N3-dCas9-DD-BirA was performed in several steps, as follows. The cDNA for dCas9 was initially cloned from EF1-dCas9-KRAS (pHAGE EF1α dCas9-KRAB was a gift from Rene Maehr and Scot Wolfe, Addgene plasmid #50919) (Kearns et al., 2014) into pINTO-C-HF (pcDNA4/TO from Invitrogen with a C-terminus HA-Flag tag) using EcoR1 and Not1. Then, BirA cDNA with a stop codon was cloned from pCDNA3.1 BirA-GFP (a gift from Vasily Ogryzko) (Kulyyassov et al., 2011) into pInto-dCas9 using Not1 and XhoI. For insertion of the FKBP(DD), the sequence was amplified from pBMN FKBP(DD)-YFP (a gift from Thomas Wandless, Addgene plamid #31763) (Banaszynski et al., 2006) using primers designed to leave Not1 overhangs. Digestion and ligation of the Not1 site in pINTO-dCas9-(Not1)-BirA and ligation of the NotI flanked FKBP(DD) fragment gave in-frame dCas9-DD-BirA. Lastly, the dCas9-DD-BirA cDNA was moved into the pINTA-N3 vector, based on the Tet-On 3G system (Clontech) through Gibson cloning to derive pINTA-dCas9-DD-BirA for expression in KH2 ESCs. See Table S1 for cloning primers.

### CRISPR-mediated genome editing

GFP was replaced for BFP in the SpCas9-2A-GFP plasmid [pSpCas9(BB)-2A-GFP (PX458 was a gift from Feng Zhang, Addgene plasmid # 48138) (Ran et al., 2013)] by Gibson cloning and made SpCas9-2A-BFP (see Table S1 for cloning primers). Then, an optimal gRNA target sequence closest to the genomic target site was chosen using the http://crispr.mit.edu design tool. The individual gRNAs (Table S2) were cloned into the SpCas9-2A-BFP plasmid via BbsI digestion and insertion (Ran et al., 2013). For knock-in of Flag-BAP, ~750 bp gBlock design were ordered from IDT or Genscript, that included 1) a homology arm corresponding to the ~350-450 bp of the H3 promoter area, 2) the Flag-BAP sequence placed after the start codon (Flag-BAP sequence: GACTACAAAGACGATGACGACAAGGGCCTGACAAGAATCCTGGAAGCTCAGAAGATCGTGAGAGGAGGCCTCGAG), and 3) a homology arm corresponding to ~200-300 bp of the H3 coding sequence. The PAM sequences of the gBlocks were mutated for correct Cas9 digestion of genomic DNA within cells (see Table S2 for primers corresponding to gBlock amplification). ESCs were then transfected with 0.5 µg of the Cas9-gRNA-BFP plasmid and 0.5 µg of amplified gBlock in Lipofectamine 2000 containing 2i media. After transfections, the medium to high BFP populations were FACS sorted and seeded at 20,000 cells per 15 cm plate. 7-10 days later, single ESC clones were selected and plated onto individual wells of a 96-well plate for genotyping. Genomic DNA was harvested via QuickExtract (Epicentre) DNA extraction, and genotyping PCRs were performed using primers surrounding the target site (Table S2), in addition to screening against Cas9 insertions with the genotyping primers FWD: ACCCAGAGAAAGTTCGACAATC and REV: GTGGTGGTAGTTGTTGATCTCG. The PCR products of the Flag-BAP positive clones were purified and sequenced to verify the presence and correct sequence of a Flag-BAP-H3 insertion.

For gRNA recruitment of dCa9-DD-BirA to 5 kb of a candidate locus in ESCs containing Flag-BAP-H3 chromatin, individual gRNAs were inserted into the pKLV-U6gRNA-EF(BbsI)-PGKpuro2ABFP (a gift from Hiroshi Ochiai, Addgene plasmid #62348) (Ochiai et al., 2015) (Table S3). Lentiviruses for each gRNA were generated in 293T cells using the packaging plasmids pMD2.G and psPAX2. Briefly, 300K 293T cells were seeded per well in a 6-well plate the day prior to transfections. For transfections, 1 µg of pAX/MD2g/gRNA vector (0.375 µg/0.125 µg/0.5 µg) was added to Lipofectamine 2000 media for overnight incubation. Fresh media was then added and 48 hr viral supernatants were collected and frozen at -80° C. For KH2 ESCs transductions, 5000 cells were seeded in 100 µl of 2i media for a 96-well plate. The following morning, tandem viral transductions were done with Polybrene containing 10 ul of each viral gRNA supernatant corresponding to ~1-16 gRNAs (total volume 160 ul) and spinoculated 2000 rpm for 90 min at 32° C. After a 6 hr incubation in a temperature (37°C)-and CO_2_-controlled chamber, the transductions were repeated with 10 µl of viral supernatants from ~17-35 gRNAs (total volume of 180 µl). Fresh 2i media was added and cells were allowed to recover for 3-5 days before FACS sorting a BFP positive population. This method allowed for a 90-95% efficiency of ES cell transduction, with ~35 gRNAs targeting a 5 kb area of a candidate locus.

### ChIP-qPCR and ChIP-seq

For native MNase chromatin preparation, nuclei were harvested from ESCs by hypotonic lysis TMSD buffer (40 mM Tris, 5 mM MgCl_2_, 0.25 M Sucrose with protease inhibitors), and pelleted for 10 min at 3600 rpm at 4° C. Nuclei were resuspended in NIB-250 (15 mM Tris-HCl, pH 7.5, 60 mM KCl, 15 mM NaCl, 5 mM MgCl_2_, 1 mM CaCl_2_, 250 mM Sucrose with protease inhibitors) containing 0.3% NP-40. A chromosome pellet was isolated by centrifuging the solubilized nuclei for 5 min at 600 rcf at 4° C. The chromosome pellet was washed with NIB-250 buffer, then resuspended with MNase digestion buffer (10 mM Hepes, 50 mM NaCl, 5 mM MgCl_2_, 5 mM CaCl_2_ with protease inhibitors) and treated with MNase (Sigma) until a DNA fragment size of 150-300 bp (1-2 nucleosomes) was attained. MNase digestion was stopped with 15 µl of EGTA (0.5 M) per 1 ml pellet and then spun at 600 rcf for 5 min at 4° C. The supernatant was placed in a fresh tube and the chromatin pellet was further processed by adding the same volume of BC500 (40 mM Tris, 5 mM MgCl_2_, 500 mM NaCl and 5% glycerol, and protease inhibitors) and EGTA (0.5 M) as in the MNase buffer plus EGTA step. The soluble chromosomes in BC500 and EGTA were then incubated for 30 min while rotating at 4° C, allowing for further enrichment of desired chromatin. Samples were pelleted at 600 rcf and the BC500 supernatant was pooled with an equal volume of MNase-treated supernatant to acquire the starting chromatin material for chromatin immunoprecipitations (IPs).

For native Flag, H3K27me3, or biotin chromatin IPs, dynabeads protein G (Invitrogen) were pre-blocked with PBS and 0.3% BSA for 1 hr, washed 5x with TE, and used to pre-clear the prepared MNase-treated chromatin for 1 hr at 4° C. For IP set up, 100 µg of pre-cleared chromatin was used with 10 µg of biotin antibody (Bethyl A150-109A), 4 µg of H3K27me3 (Cell Signaling C36B11), or FLAG-M2 agarose beads (Sigma) and IP was performed overnight with slow rotation at 4° C. For quantitative analysis of ChIPs, 0.5 µg of *Drosophila S2* chromatin was spiked-in to 100 µg of sample (1:200 concentration). The following morning, pre-blocked dynabeads protein G were added to the IPs and incubated for 2-3 hr at 5 µl slurry/1 µg antibody with 0.2 µl of *Drosophila*-specific H2Av antibody for spike-in control in each sample. IPs were then washed three times with 1 ml BC300 buffer (40 mM Tris, 5 mM MgCl2, 300 mM NaCl and 5% glycerol with protease inhibitors), once with 1 ml BC100 buffer (40 mM Tris, 5 mM MgCl2, 100 NaCl and 5% glycerol with protease inhibitors), and a quick wash with 1 ml TE + 50 mM NaCl. Except for Flag-ChIPs, beads were then resuspended in 125 µl of TE, 3 µl of 10% SDS (TES) and incubated at 65° C for 1 hr. For Flag-ChIPs, the bound complexes were eluted with 0.25 µg FLAG peptide (Sigma) in 100 µl BC100 buffer overnight with slow rotation at 4° C. The Flag-ChIPs and TES-treated chromatin were then removed from their respective beads and the input was thawed and resuspended in 125 µl TES. Input and samples were then digested for 2-4 hr with 8 µg of proteinase K at 55° C while shaking. All samples were PCR column purified, eluted in 50 µl, and diluted 1:4 with water for further qPCR studies. For qPCR quantification, 5 µl SYBR Green I Master mix (Roche), ROX reference dye, 1 µl 5 µM primer pair, 4 µl-diluted DNA were mixed for PCR amplification, and detected by a Stratagene Mx3005p instrument. The data was then quantified and described in the corresponding figure legends. For *Drosophila S2* chromatin, primers were FWD: TGGCTAGACTTTTGCGTCCT and REV: TACCAAAAGCCGTCCAAATC and other qPCR primers are listed in Table S4.

For native Flag, H3K27me3, or biotin ChIP-Seq, DNA was eluted in 30 µl elution buffer and libraries for ChIP-seq were prepared according to the manufacturer’s instructions (Illumina). Briefly, IP’ed DNA (~1-30 ng) was end-repaired using End-It Repair Kit (Epicenter), tailed with deoxyadenine using Klenow exo-(New England Biolabs), and ligated to custom adapters with T4 Rapid DNA Ligase (Enzymatics). Fragments of 200-400 bp were size-selected using Agencourt AMPure XP beads, and subjected to PCR amplification using Q5 DNA polymerase (New England Biolabs). Libraries were quantified by qPCR using primers annealing to the adapter sequence and sequenced on an Illumina HiSeq. Barcodes were utilized for multiplexing.

Fixed chromatin IP experiments were described previously (Gao et al., 2012). Briefly, cells were fixed with 1% Formaldehyde. Nuclei were isolated using buffers in the following order: LB1 (50 mM HEPES, pH 7.5 at 4° C, 140 mM NaCl, 1 mM EDTA, 10% Glycerol, 0.5% NP40, 0.25% Triton X; 10 min at 4° C), LB2 (10 mM Tris, pH 8 at 4° C, 200 mM NaCl, 1 mM EDTA, 0.5 mM EGTA; 10 min at RT), and LB3 (10 mM Tris, pH 7.5 at 4° C, 1 mM EDTA, 0.5 mM EGTA, and 0.5% N-Lauroylsarcosine sodium salt). Chromatin was fragmented to an average size of 250 bp using a Diagenode Bioruptor. For fixed chromatin, Cas9 antibody from Millipore (MAC133-clone 7A9) was used.

### RNA Purification and Quantitative PCR

Total RNA extractions were performed using Roche High pure RNA isolation kit. Superscript III Reverse Transcription Reagents (Invitrogen) and random hexamers were used to prepare cDNAs. For qPCR quantification, 5 µl SYBR Green I Master mix (Roche), ROX reference dye, 1 µl IDT PrimeTime Primer set for corresponding assays, 1 µl cDNA were mixed for PCR amplification, and detected by a Stratagene Mx3005p instrument. For quantitative analysis, *Gata2* and *Gata6* expression was normalized to *Actb* expression.

## QUANTIFICATION AND STATISTICAL ANALYSIS

### ChIP-seq data analysis

ChIP-seq were analyzed as previously reported (Stafford et al., 2018). Briefly, sequence reads were mapped with Bowtie (Langmead et al., 2009) and Spike-in normalization was achieved using *Drosophila Melanogaster* DNA aligned to Illumina igenomes dm3 (Orlando et al., 2014). Endogenous and exogenous scaling factors were computed from the bam files (1/(number_mapped_reads/1000000)). Endogenous scaling factors were applied to the data before input subtraction (without scaling). The RPM normalization was then reversed before scaling with exogenous factors. The scripts used to perform spike-in scaling have been integrated in a bioconductor package ChIPseqSpike (https://bioconductor.org/packages/3.7/bioc/html/ChIPSeqSpike.html).

### ChIP-qPCR and statistical significance

Native biotin enrichment levels were normalized to 5% input followed by *Drosophila* chromatin spike-in levels. For time course experiments, data was minus-Dox (-Dox) control subtracted and error bars represent standard deviation of three biological replicates. GraphPad Prism 7.0 was used for statistical analysis (2way ANOVA). A *p* value ≤ 0.05 was considered statistically significant with *** denotes *p*<0.001 and **** denotes *p*<0.0001.

## DATA AND SOFTWARE AVAILABILITY

The ChIP-seq data has been deposited in the Gene Expression Omnibus (GEO) under GSE120056 and will be made available immediately upon publication.

---

**Figure S1.**
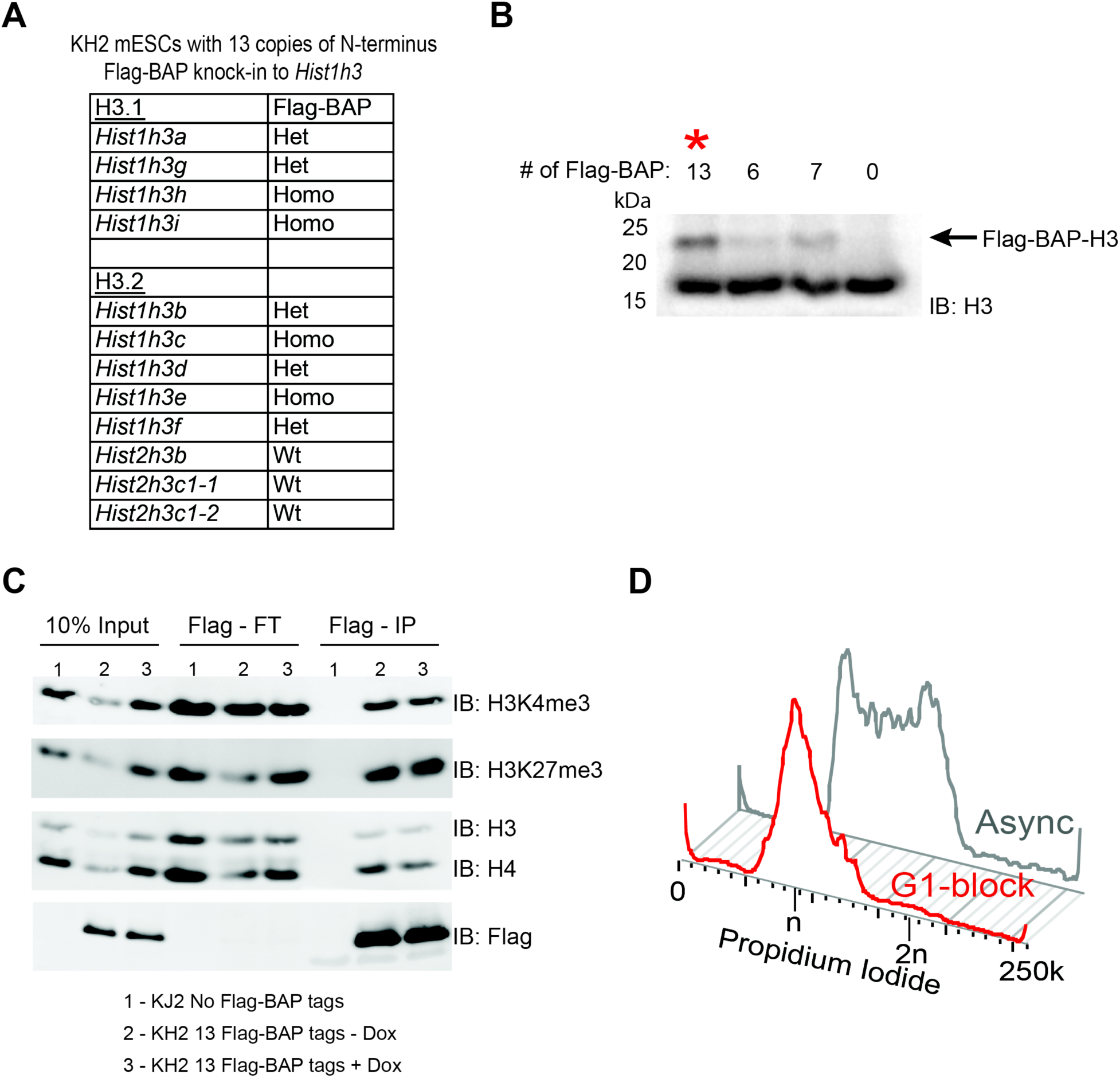
Endogenous tagging of H3.1 and H3.2 histones with Flag - Biotin Acceptor Peptide (BAP) in KH2 mESCs. Related to Figure 1. (**A**) Overview and genotype of KH2 mESC lines containing 13 copies of Flag-BAP knock-ins to H3.1 and H3.2 loci. (**B**) Western blot analysis for histone H3. The upper band corresponds to the endogenous number of Flag-BAP-H3 copies in the cell and the red asterisk denotes the cell line used for further studies. (**C**) Flag IPs and flow-through (FT) of KH2 cells containing Flag-BAP tags and controls. Analysis highlights the pull-down of H3K4me3 and H3K27me3 chromatin. (**D**) Flow cytometry analysis showing DNA content of mESCs following double thymidine G1 synchronization.

**Figure S2.**
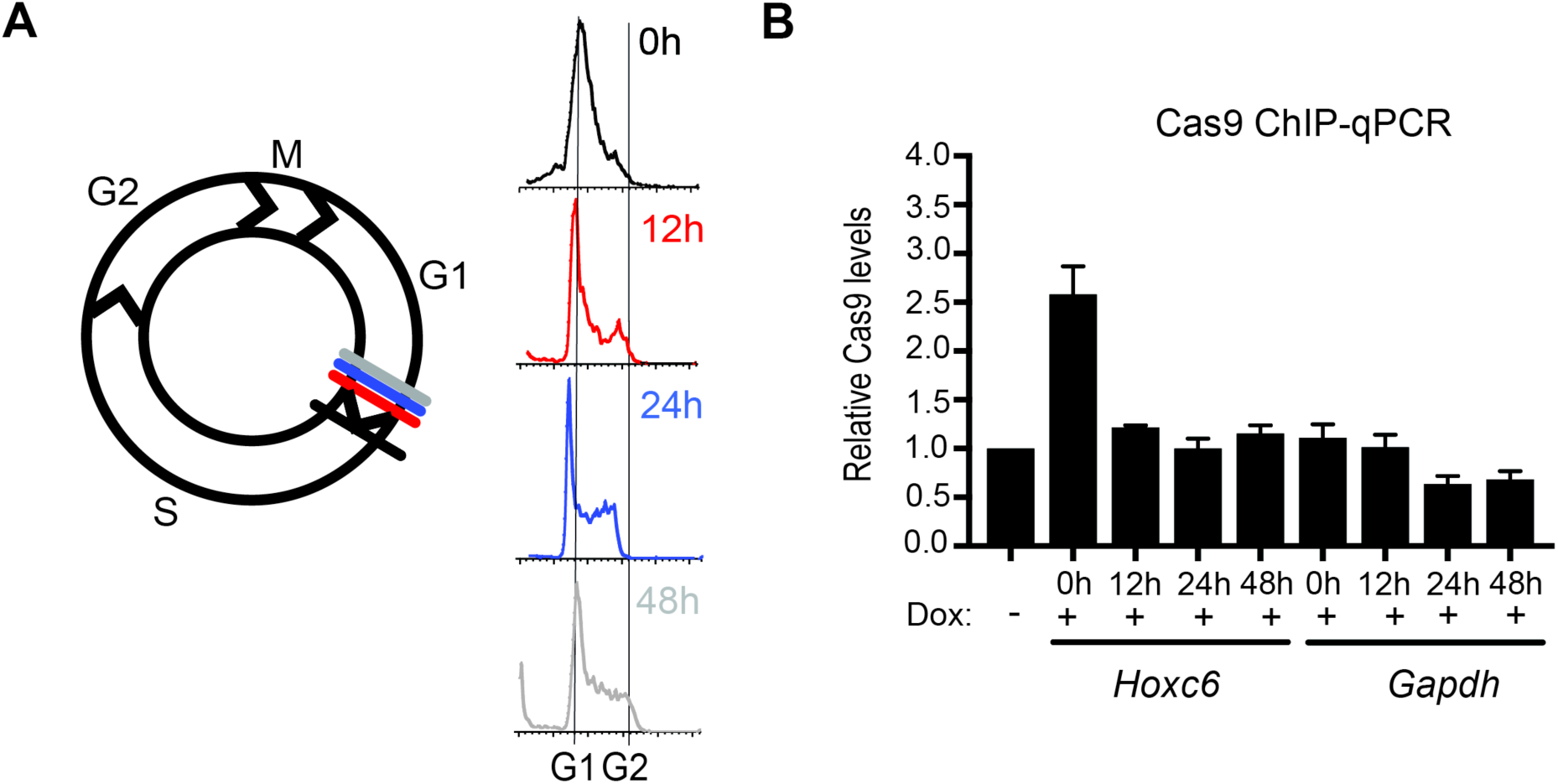
Inducibility of dCas9-DD-BirA during cell division in mESCs. Related to Figure 1. (**A**) Cell cycle analysis corresponding to G1/S-blocked and released mESCs at time 0 hr -1 cell, 12 hr -2 cell, 24 hr - 4 cells, and 48 hr - 16 cells. (**B**) Cas9 ChIP-qPCR analysis of G1/S-blocked and released mESCs following a 6 hr pulse with minimal doxycycline (Dox) and exogenous biotin in cells targeting dCas9-DD-BirA to the *Hoxc6* locus compared to *Gapdh* controls. Data was normalized to 5% input and error bars represent standard error of three biological replicates.

**Figure S3.**
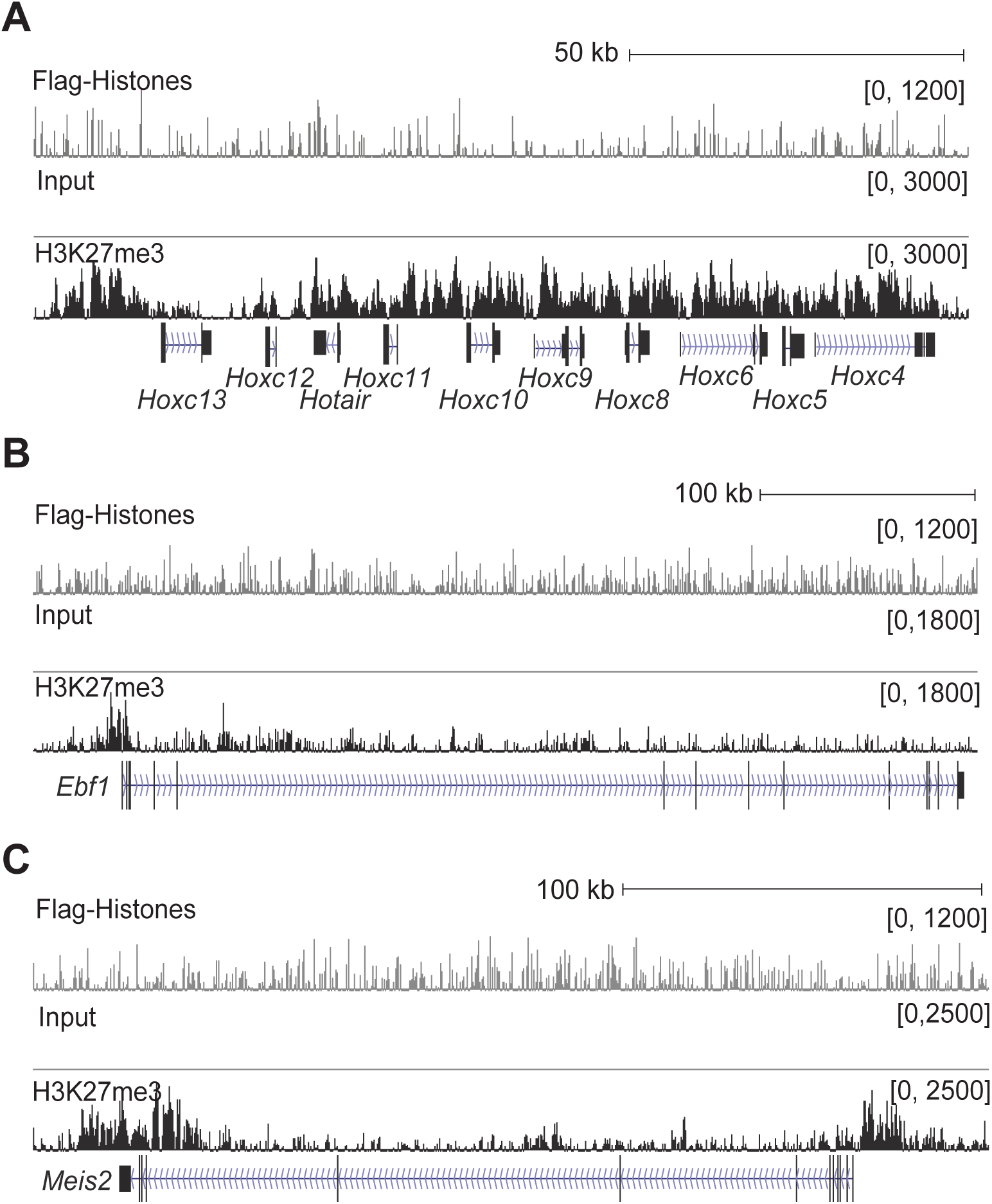
Candidate repressed chromatin domains in ESCs. Related to Figure 2. (**A-C**) Native Flag and H3K27me3 ChIP-seq analysis of G1/S-blocked cells at the *Hoxc* cluster (**A**), *Ebf1* (**B**), and *Meis2* (**C**) candidate loci.

**Figure S4.**
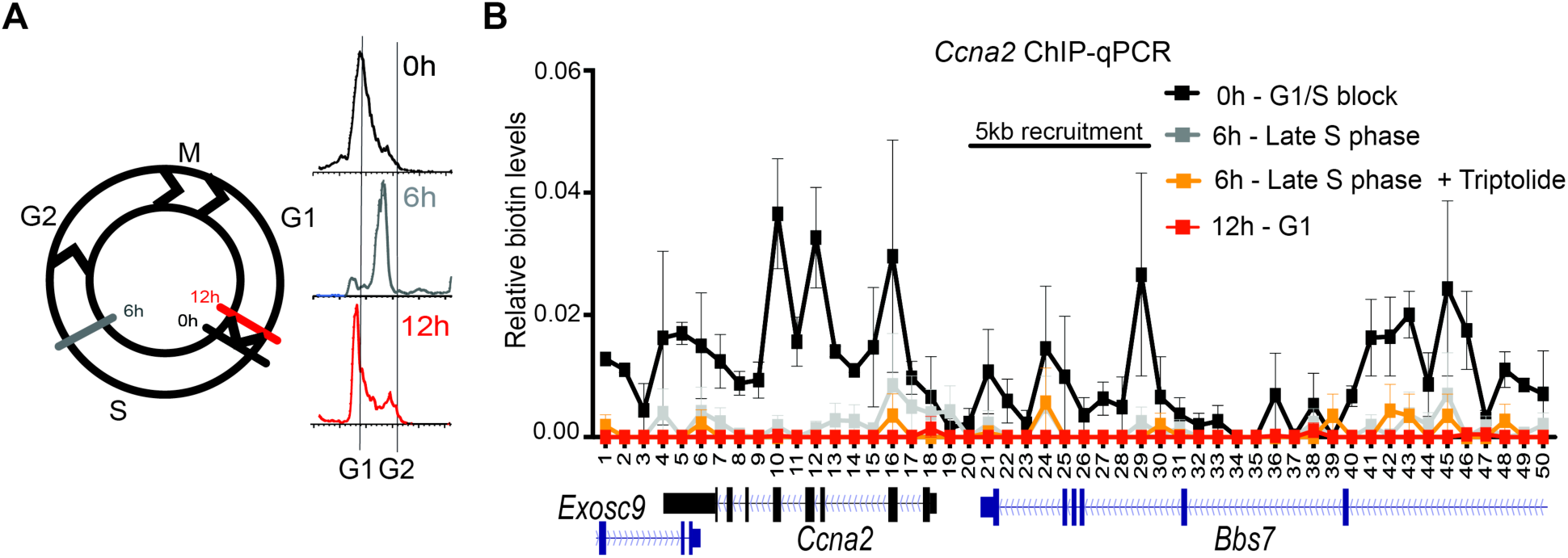
Biotin-H3 in *Ccna2* active chromatin is dynamic during S-phase progression. Related to Figure 3. (**A**) Cell cycle analysis corresponding to G1/S-blocked and released mESCs at time 0 hr - 1 cell, 6 hr -1 cell (late S-phase), and 12 hr - 2 cells. (**B**) Native biotin ChIP-qPCR from G1/S-blocked and released mESCs following a 6 hr pulse of minimal Dox and exogenous biotin in cells targeting BirA to the *Ccna2* locus. The time course represents time at which cells were released into S-phase with or without 1 mM Triptolide: 0 hr -1 cell, 6 hr - Late S phase, and 12 hr - 2 cells. Data is the average of two biological replicates spanning a 25 kb area at a resolution of 500 bp. All biotin enrichment levels are relative to input and normalized to *Drosophila* chromatin spike-in followed by subtraction of the -Dox control. Error bars represent standard error of three biological replicates.

**Figure S5.**
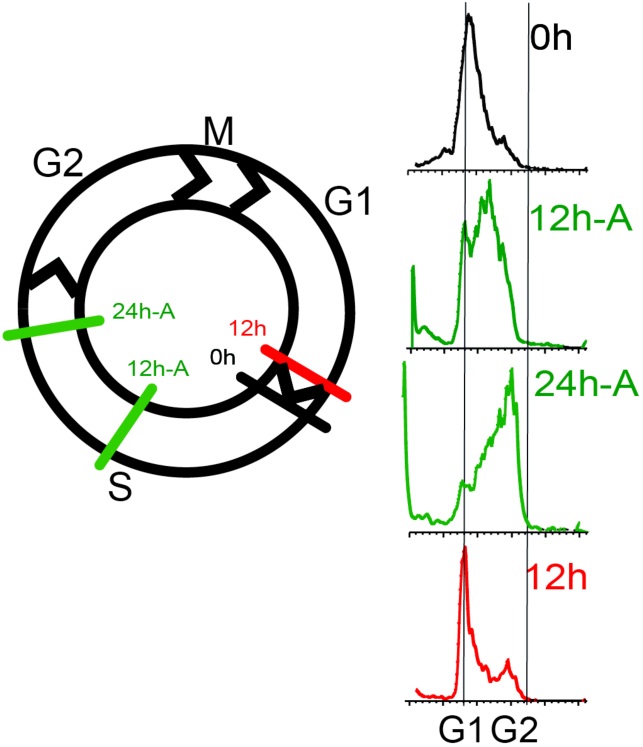
Aphidicolin treatment sustains ESCs in S-phase. Related to Figure 4. Cell cycle analysis corresponding to G1/S-blocked and released ESCs treated with 1 mg/ml Aphidicolin for 12 hr or 24 hr to sustain cells in S-phase.

**Figure S6.**
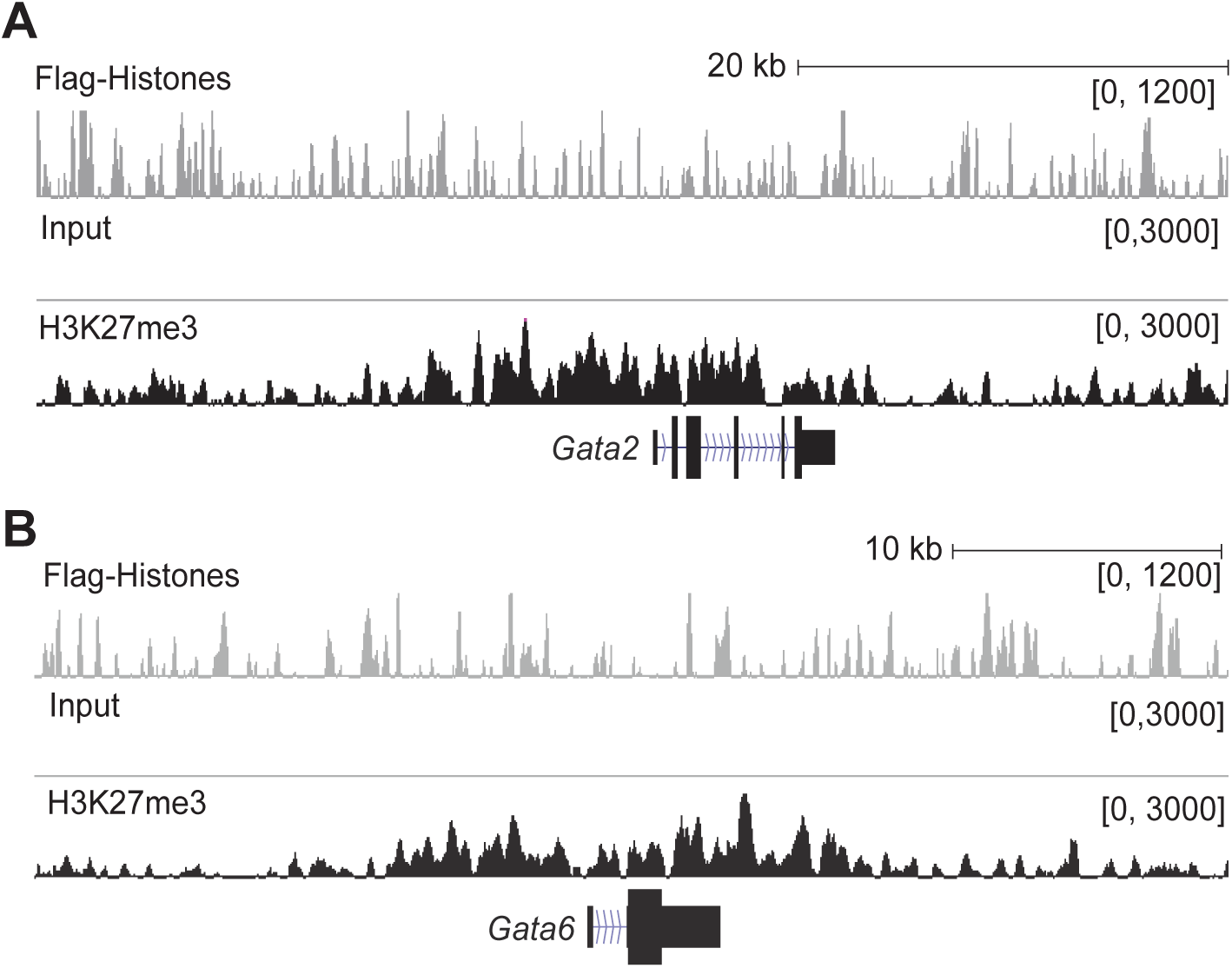
*Gata* factors are repressed in ESCs. Related to Figure 5. (**A-B**) Native Flag and H3K27me3 ChIP-seq analysis of G1/S-blocked cells at the *Gata2* (**A**) and *Gata6* (**B**) candidate loci.

